## Supplementary material for "Dynamic WT1 expression during gastrulation specifies peritoneal smooth muscle fate independently from mesothelial fate": all supplementary information

**Supplementary Table 1. List of litters and embryos analysed by Tamoxifen-induced lineage tracing until just before birth.**

| Stage at Tam dose | Stage at analysis | Lineage marker | Number of embryos analysed | mesentery, mid-intestine | | | stomach | | | colon | | |
|---|---|---|---|---|---|---|---|---|---|---|---|---|
| | | | | mesothelium | vSMCs | viSMCs | mesothelium | vSMCs | viSMCs | mesothelium | vSMCs | viSMCs |
| E14.5 | E19.5 | LacZ | 3 | +++ | | | ++ | | | ++ | | |
| | E18.5 | LacZ | 9 | +++ | | | | | | | | |
| E13.5 | E17.5 | LacZ | 3 | ++++ | | | ++++ | | | ++++ | | |
| E12.5 | E19.5 | LacZ | 1 | ++++ | | | | | | | | |
| E11.5 | E17.5 | LacZ | 3 | +++ | | | ++ | | + | | | |
| | E19.5 | LacZ | 6 | ++ | +* | | | | | | | |
| | E19.5 | mTmG | 2 | +++ | | | | | | | | |
| E10.5 | E18.5 | LacZ | 9 | ++ | | | + | | + | + | | + |
| E9.5 | E19.5 | LacZ | 2 | ++ | + | | | | | | | |
| | E17.5 | LacZ | 1 | ++ | | | ++ | | | + | | ++ |
| E8.5 | E18.5 | LacZ | 2 | + | + | + | | | | | | |
| | E18.5 | LacZ | 2 | + | | + | + | | + | | | ++ |
| | E17.5 | mTmG | 6 | + | + | + | | | | | | |
| E7.5 | E19.5 | LacZ | 2 | | ++ | | | | | | | + |
| | E18.5 | LacZ | 4 | | ++ | + | | | + | | | + |
| | E18.5 | mTmG | 10 | | ++ | + | | | | | | |

Greyed fields indicate that no analysis was performed.

* indicates that 2 embryos of 6 with Tamoxifen at E11.5, had labelled viSMCs.

+ a few sparse mesothelial cells in mesentery

++ some patches of mesothelial cells

+++ mesothelial cells, not a full coverage

++++ full mesothelial coverage

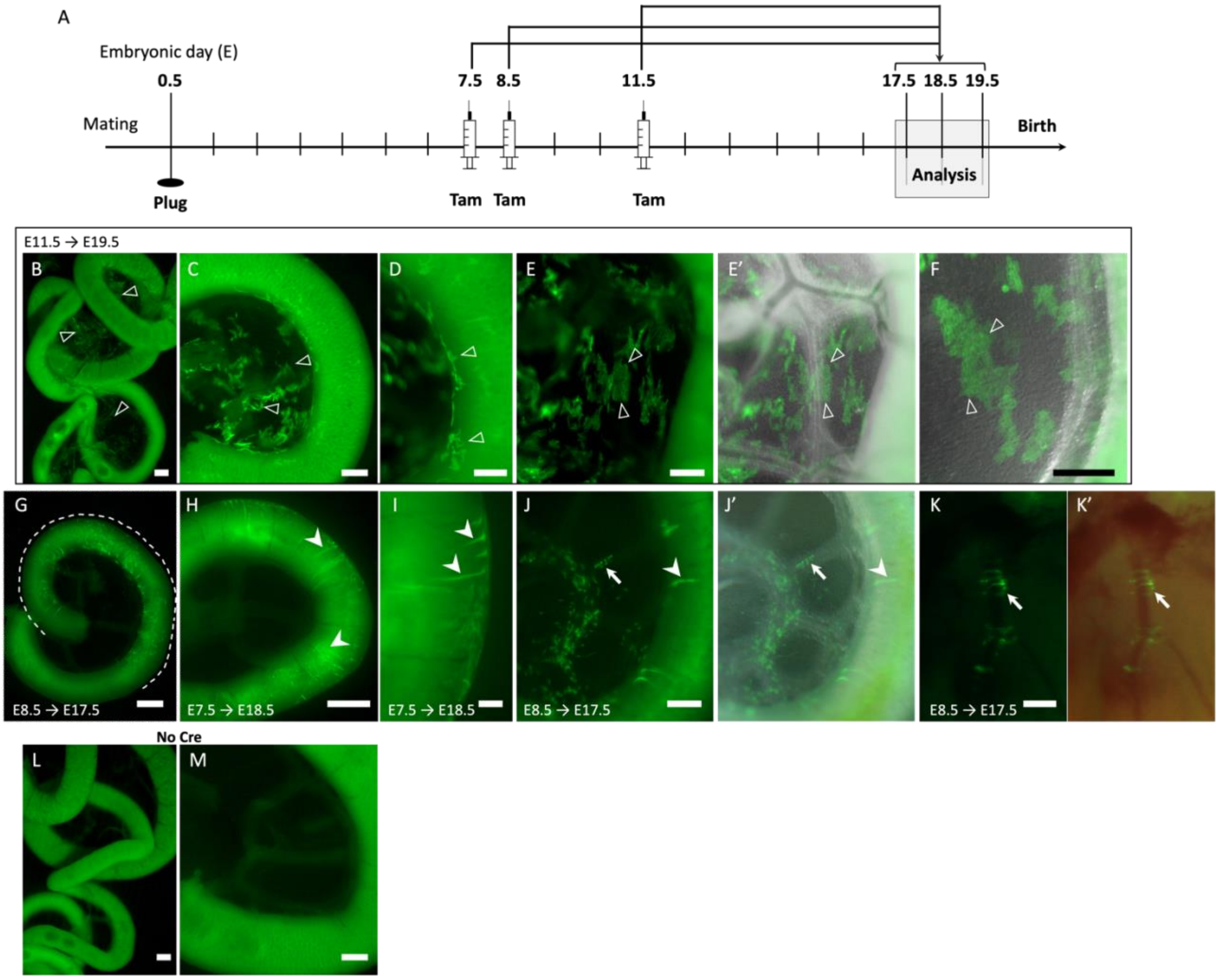


**Supplementary Figure 1. Wt1 lineage in embryos using the *Rosa*$^{mTmG}$ reporter.**

**A.** Timeline of experiment: analysis of embryos after time mating of *Wt1*$^{CreERT2/+}$;*Rosa26*$^{mTmG/mTmG}$ males with CD1 females and Tam dosing at E11.5, E8.5 or E7.5.

**B-F.** After Tam dosing at E11.5 and analysis at E19.5, GFP-expressing cells were observed within the mesentery (open arrowheads) along the entire lengths of the small intestine. This included patches of GFP-positive stained cells (open arrowheads) over the intestine (D), and across the mesentery (C, E, E', F).

**G-K'.** When Tam was given at E7.5 or E8.5 and embryos analysed at E17.5 or E18.5, GFP-expressing cells were found in restricted segments (G, stippled line) along the small intestine only, and comprised circular visceral smooth muscle cells (arrowheads, H, I, J, J'). GFP expression was also found in cells surrounding mesenteric vessels (arrows, J, J', K, K').

**L, M.** In pups negative for the CreERT2 modification and after Tam at E11.5 and analysis at E19.5, no GFP-positive cells were observed.

Tam, Tamoxifen. Scale bars, 500 µm (B, G, H, L), 300 µm (C, M), 200 µm (D, E, E', F, I, J, J', K, K').

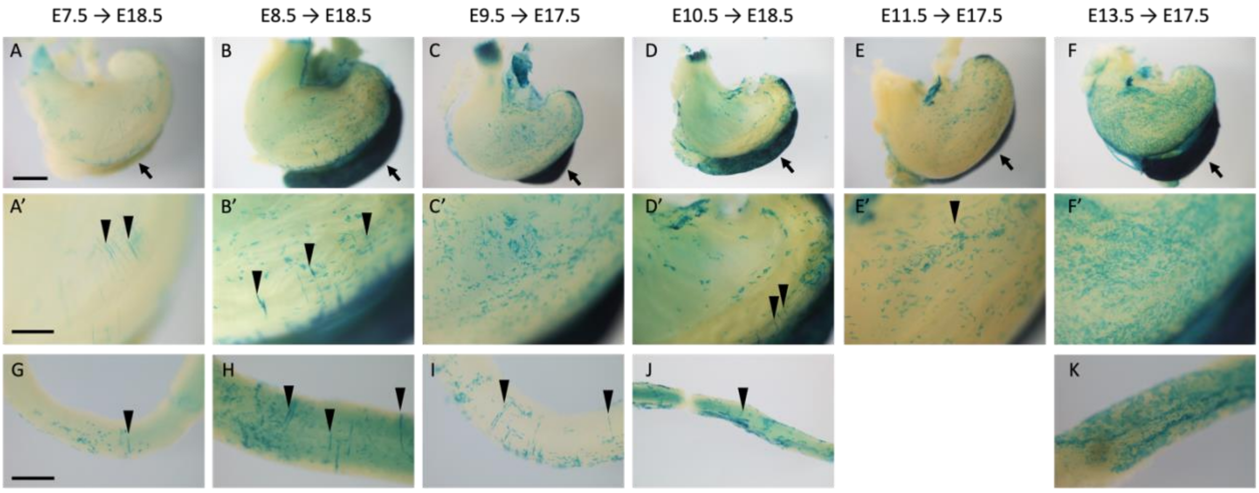


**Supplementary Figure 2. Embryonic contribution of cells of the *Wt1*-lineage to visceral mesothelium or visceral and vascular smooth muscle in colon, stomach and spleen.** Schematic of the study design as in Figure 1A; CD1 dams were time mated with *Wt1*$^{CreERT2/+}$;*Rosa26*$^{LacZ/LacZ}$ males and received Tam at the indicated embryonic stages; embryos were dissected between E17.5 and E19.5, and XGal stained. Representative whole mount-stained organs: A-F' show stomach with spleen attached (arrow), and G-K show colon segments.

**A, A'.** When Tam was given at E7.5 and embryos analysed at E18.5, there were visceral smooth muscle cells (arrowheads) labelled in the stomach.

**B, B', D, D', E, E'.** When Tam had been given at E8.5 (E18.5), E10.5 (E18.5) or E11.5 (E17.5), LacZ-positive cells consisted of both mesothelial and visceral smooth muscle cells (arrowheads).

**C, C'.** Interestingly, Tam dosing at E9.5 and analysis at E17.5 resulted in no labelled visceral smooth muscle cells, only mesothelial cells covering the stomach.

**F, F'.** Tam dosing at E12.5 and analysis at E19.5 also showed only labelled mesothelial cells which covered the stomach almost completely. The spleen (arrow, A-F) was labelled only in embryos when Tam had been given at E8.5 or later stages.

**G-K.** When Tam was given at E7.5, E8.5, E9.5 or E10.5, there were labelled visceral smooth muscle cells in the colon. Labelled mesothelial cells were only found in embryos that had been given Tam at E9.5 or later.

Images of stomach or colon from embryos given Tam at E14.5 not shown.

Scale bars, 500 µm (B, C, D, F, H), 200 µm (E, G, I, J).

A

| Stage | Sample Size | Total No. of Cells | Average No. of Cells | Average No. of *Wt1* Cells | % of *Wt1*+ Cells |
|---|---|---|---|---|---|
| **E6.5** | 3 | 3697 | 1232 | 20 | 1.60 |
| **E6.75** | 1 | 2169 | 2169 | 35 | 1.61 |
| **E7.0** | 6 | 16571 | 2762 | 46 | 1.66 |
| **E7.25** | 3 | 15294 | 5098 | 33 | 0.65 |
| **E7.5** | 5 | 12876 | 2575 | 23 | 0.91 |
| **E7.75** | 4 | 17720 | 4430 | 32 | 0.72 |
| **E8.0** | 4 | 22059 | 5515 | 45 | 0.82 |
| **E8.25** | 3 | 18642 | 6214 | 53 | 0.85 |
| **E8.5** | 4 | 20978 | 5245 | 59 | 1.12 |

B

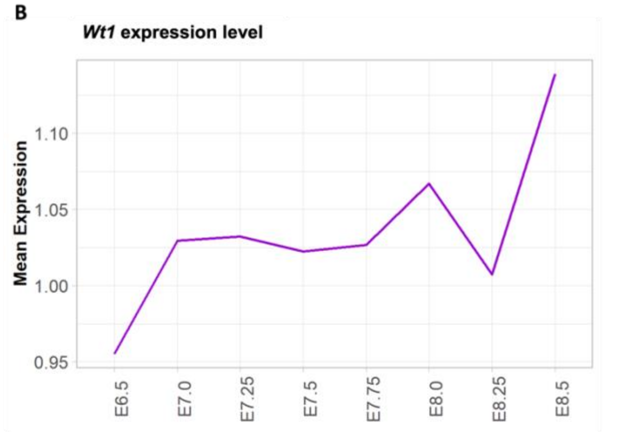


C

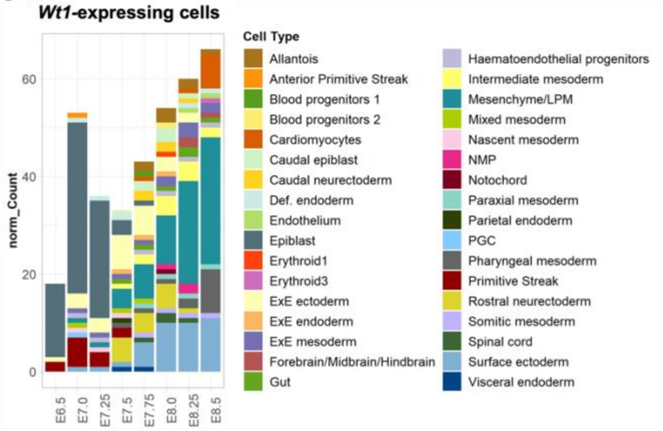


D

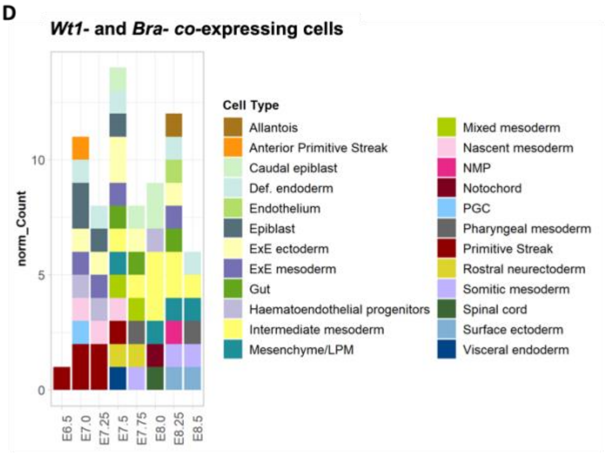


E

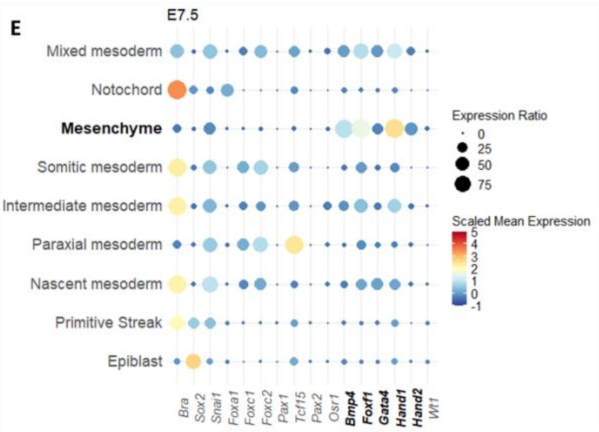


F

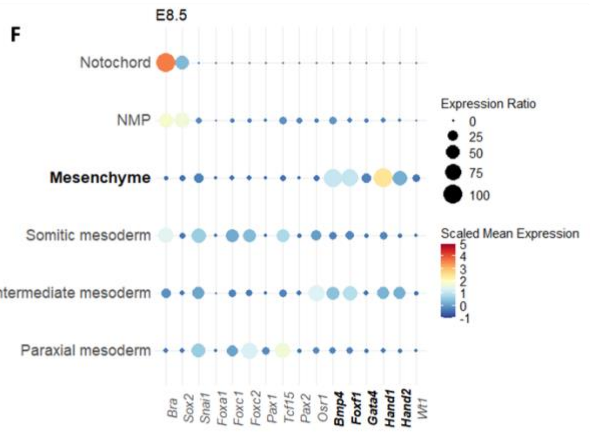


G

| Group | Markers |
|---|---|
| SMC_geneSet | *Ppp1r12b, Lmod1, Myl9, Tpm2, Actg2, Cnn1, Tagln, Myh11, Mylk, Acta2, Flna, Mrtfa, Mrtfb, Pdgfrb, Bmp4* |
| MC_geneSet | *Aldh1a2, Prrx2,Rspo1, Upk3b, Wnt2b, Msln, Fgf18, Muc16, Lrrn4, Meis3, Gata5, Gata6, Tcf21, Tbx18, Pdpn* |

H

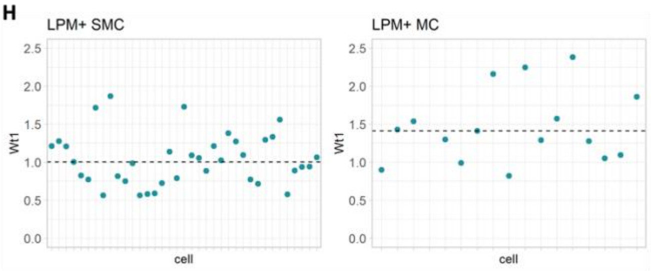


I

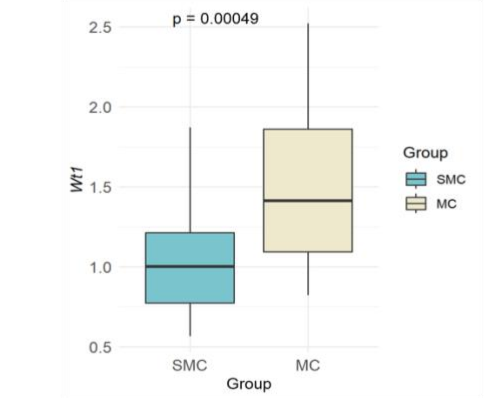


J

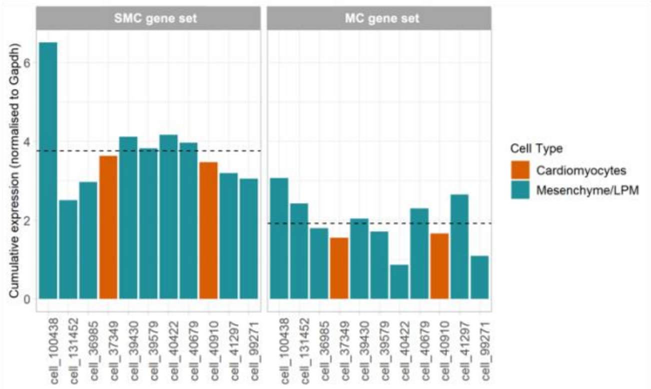

**Supplementary Figure 3.1. Distribution of *Wt1*-expressing cells in gastrulation stage embryos.**

**A.** Total number of cells with *Wt1* expression greater than zero in each stage. E6.75 was excluded from downstream analysis due to the small sample size (n=1).

**B.** Expression levels of *Wt1* across gastrulation stages.

**C.** Dynamic distribution of *Wt1*-expressing cells in different tissues across embryonic stages.

**D.** Dynamic distribution of *Wt1*- and *Bra*-co-expressing cells in different tissues across embryonic stages.

**E, F.** Mean expression of mesodermal lineage marker genes at E7.5 and E8.5 to characterize the tissue termed 'mesenchyme' in the dataset. Expression ratio indicates percentage number of gene-expressing cells in comparison to total number of cells of each cell type. In 'mesenchyme', about 75% of cells have elevated expression of the lateral plate mesoderm (LPM) markers *Bmp4*, *Foxf1* and *Hand1* at E7.5 and E8.5. *Hand2* expression was slightly increased in E8.5 compared to E7.5 and indicates an LPM-derived cell type.

**G.** List of gene markers used for SMC and mesothelial cell (MC) signature scoring.

**H.** Cumulative expression of signature markers in the 11 cells that scored above threshold in *Wt1*-expressing cells with both SMC and MC signature at E8.5. Dashed lines represent mean expression. X-axis lists individual cell identities, and cumulative expression (y-axis) is normalized to *Gapdh* expression.

**I.** *Wt1* expression levels of LPM cells with SMC or MC characteristics at E8.5. LPM cells with SMC characteristics had significantly lower *Wt1* expression than those with MC signature.

**J.** Cumulative expression of all marker genes of the respective signatures in the 11 cells that scored above threshold in both MC and SMC signature at E8.5. Dashed lines represent mean expression.

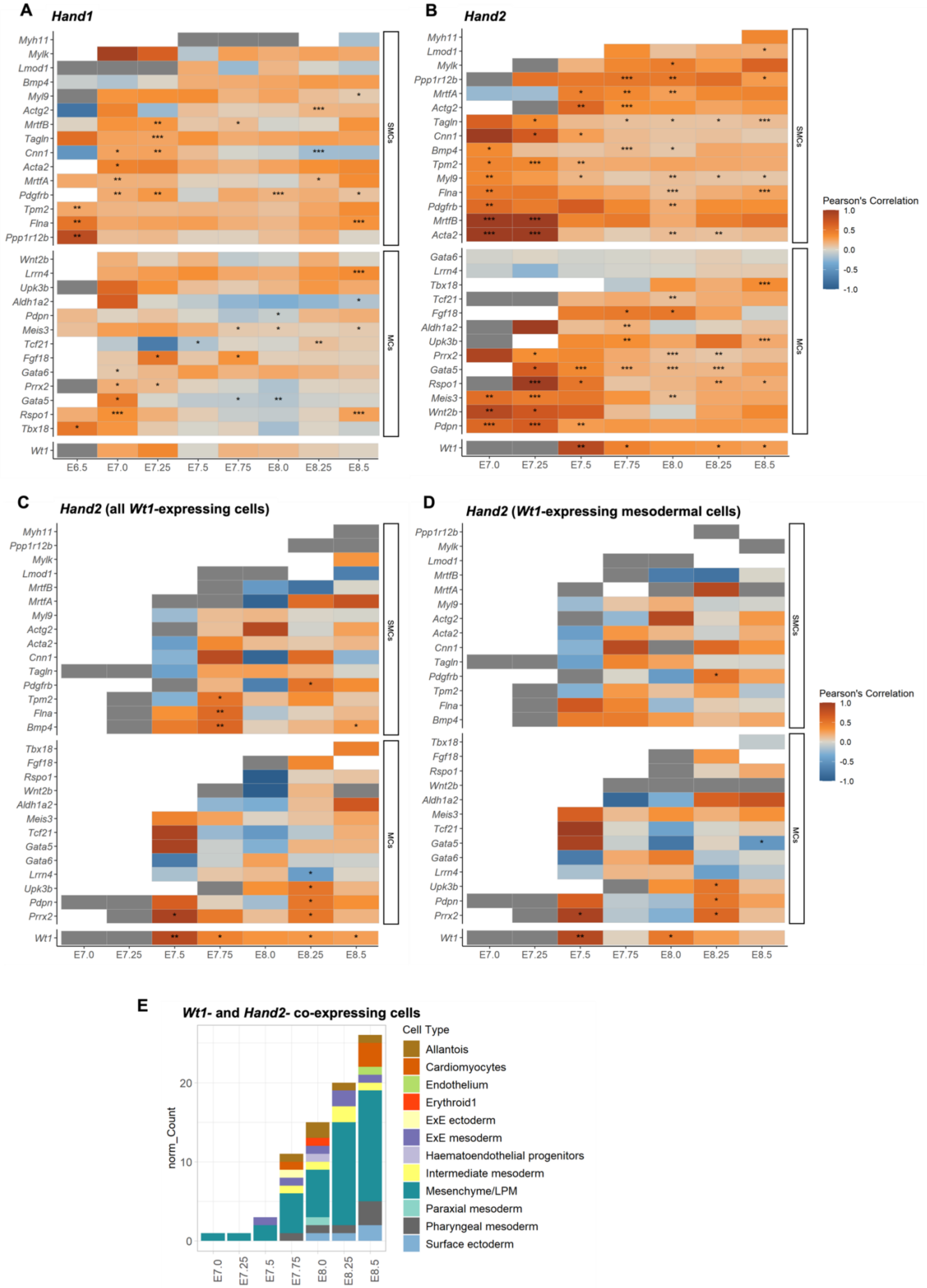
A
Hand1
Myh11
Mylk
Lmod1
Bmp4
Myl9
Actg2
MrtfB
Tagln
Cnn1
Acta2
MrtfA
Pdgfrb
Tpm2
Flna
Ppp1r12b
Wnt2b
Lrrn4
Upk3b
Aldh1a2
Pdpn
Meis3
Tcf21
Fgf18
Gata6
Prrx2
Gata5
Rspo1
Tbx18
Wt1
E6.5
E7.0
E7.25
E7.5
E7.75
E8.0
E8.25
E8.5
SMCs
MCs
B
Hand2
Myh11
Lmod1
Mylk
Ppp1r12b
MrtfA
Actg2
Tagln
Cnn1
Bmp4
Tpm2
Myl9
Flna
Pdgfrb
MrtfB
Acta2
Gata6
Lrrn4
Tbx18
Tcf21
Fgf18
Aldh1a2
Upk3b
Prrx2
Gata5
Rspo1
Meis3
Wnt2b
Pdpn
Wt1
E7.0
E7.25
E7.5
E7.75
E8.0
E8.25
E8.5
SMCs
MCs
Pearson's Correlation
1.0
0.5
0.0
-0.5
-1.0
C
Hand2 (all Wt1-expressing cells)
Myh11
Ppp1r12b
Mylk
Lmod1
MrtfB
MrtfA
Myl9
Actg2
Acta2
Cnn1
Tagln
Pdgfrb
Tpm2
Flna
Bmp4
Tbx18
Fgf18
Rspo1
Wnt2b
Aldh1a2
Meis3
Tcf21
Gata5
Gata6
Lrrn4
Upk3b
Pdpn
Prrx2
Wt1
E7.0
E7.25
E7.5
E7.75
E8.0
E8.25
E8.5
SMCs
MCs
D
Hand2 (Wt1-expressing mesodermal cells)
Ppp1r12b
Mylk
Lmod1
MrtfB
MrtfA
Myl9
Actg2
Acta2
Cnn1
Tagln
Pdgfrb
Tpm2
Flna
Bmp4
Tbx18
Fgf18
Rspo1
Wnt2b
Aldh1a2
Meis3
Tcf21
Gata5
Gata6
Lrrn4
Upk3b
Pdpn
Prrx2
Wt1
E7.0
E7.25
E7.5
E7.75
E8.0
E8.25
E8.5
SMCs
MCs
Pearson's Correlation
1.0
0.5
0.0
-0.5
-1.0
E
Wt1- and Hand2- co-expressing cells
norm_Count
0
10
20
E7.0
E7.25
E7.5
E7.75
E8.0
E8.25
E8.5
Cell Type
Allantois
Cardiomyocytes
Endothelium
Erythroid1
ExE ectoderm
ExE mesoderm
Haematoendothelial progenitors
Intermediate mesoderm
Mesenchyme/LPM
Paraxial mesoderm
Pharyngeal mesoderm
Surface ectoderm

**Supplementary Figure 3.2. *Hand1*- and *Hand2*-expressing cells correlate with SMC- and mesothelial cell signatures.**

**A, B.** Correlation plots of *Hand1* (A) *and Hand2* (B) expression against SMC marker or MC marker gene expression in total *Hand1*- or *Hand2*-expressing cells at different embryonic time points. *Hand1* expression was positively and highly significantly correlated with the expression of 3-4 SMC markers and 1-4 mesothelial cell markers at between E6.5 and E7.25. Correlation with both signatures is weaker and less significant in later embryonic stages. *Hand2* expression was positively and highly significantly correlated with the expression of 8 SMC markers and 3 mesothelial cell markers at E7.0, and was highly dynamic until E8.5, generally showing high positive correlation and significance across both signature sets. *Wt1* expression is significantly correlated with *Hand2* from E7.5 onwards but not with *Hand1*.

**C, D.** *Hand2* expression correlated less well with SMC marker or mesothelial cell marker gene expression in all cells (C) or all mesodermal cells (D). At E7.5, the most significant correlation was with *Wt1* and *Prrx2* in the mesothelial cell marker group in all cells as well as mesoderm cells, while at E7.75, correlation was significant with the SMC markers *Tpm2*, *Flna* and *Bmp4* as well as *Wt1* amongst all cells. In all cells and all mesodermal cells, *Hand2* expression correlated significantly with *Upk3b*, *Pdpn* and *Prrx2* in the mesothelial cell marker group at E8.25. White boxes indicate total absence of data at respective time points; grey boxes indicate less than three data points (p-values: ***<0.001; ** <0.01; * <0.05).

**E.** Dynamic distribution of *Wt1-, Hand2-co*-expressing cells in different tissues across embryonic stages.

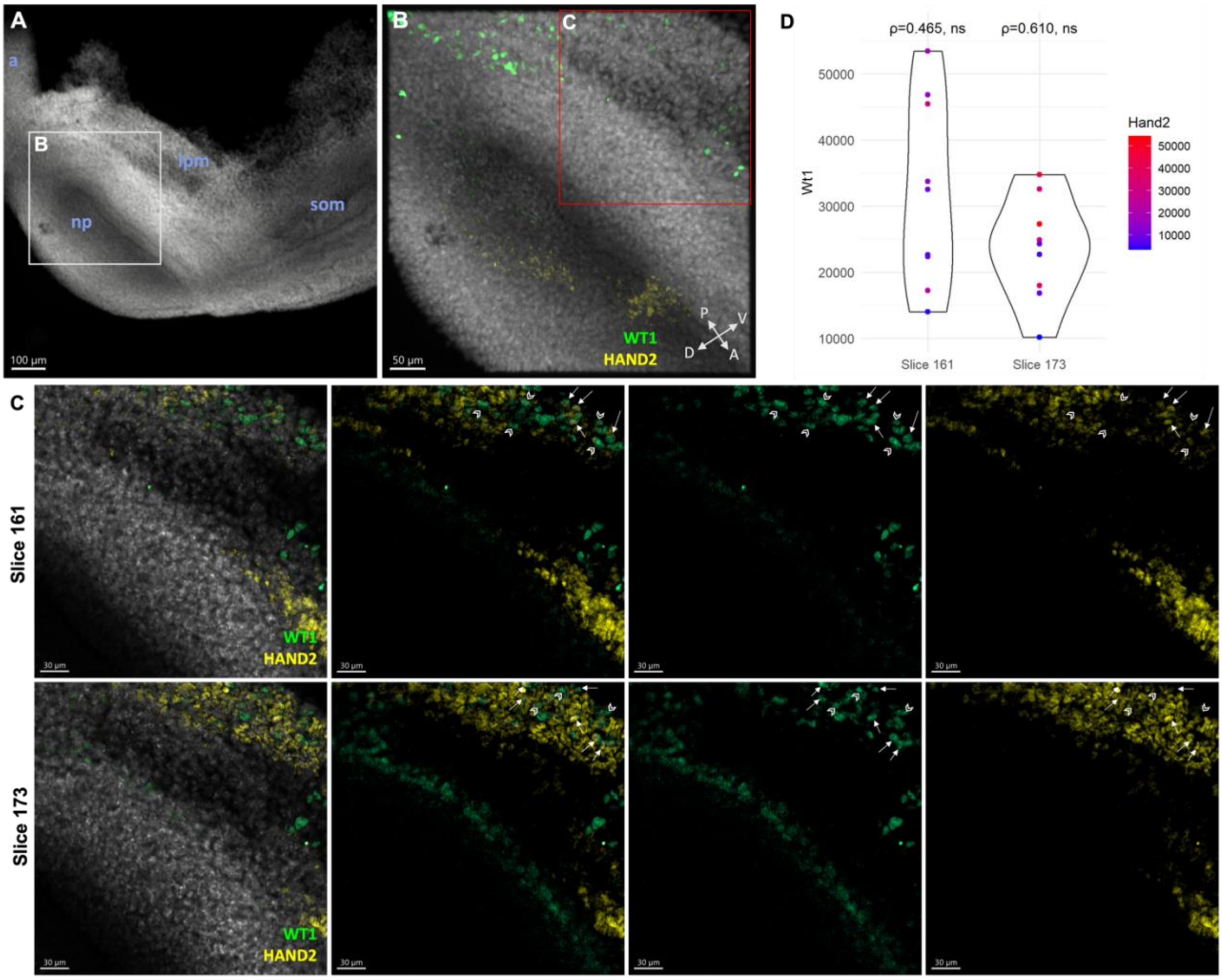


**Supplementary Figure 3.3. WT1 and HAND2 co-expression in the LPM of the E8.5 mouse embryo.**

**A, B.** Whole-mount staining of WT1 and HAND2 in the E8.5 mouse tail bud. Annotations: a, allantois, lpm, lateral plate mesoderm, np, neural plate, som, somites.

**C.** Two focal planes of a z-stack (sagittal view) showing the expression of WT1 and HAND2 in the LPM. A balanced number of WT1-positive cells co-expressing (filled arrows) and not co-expressing (hollow arrows) HAND2 were observed.

**D.** Quantification of expression levels of cells co-expressing WT1 and HAND2 at the two focal planes indicated in panel C. Spearman correlation Rho value (ρ) were calculated for each focal plane. Both slices showed p-values>0.05 (ns).

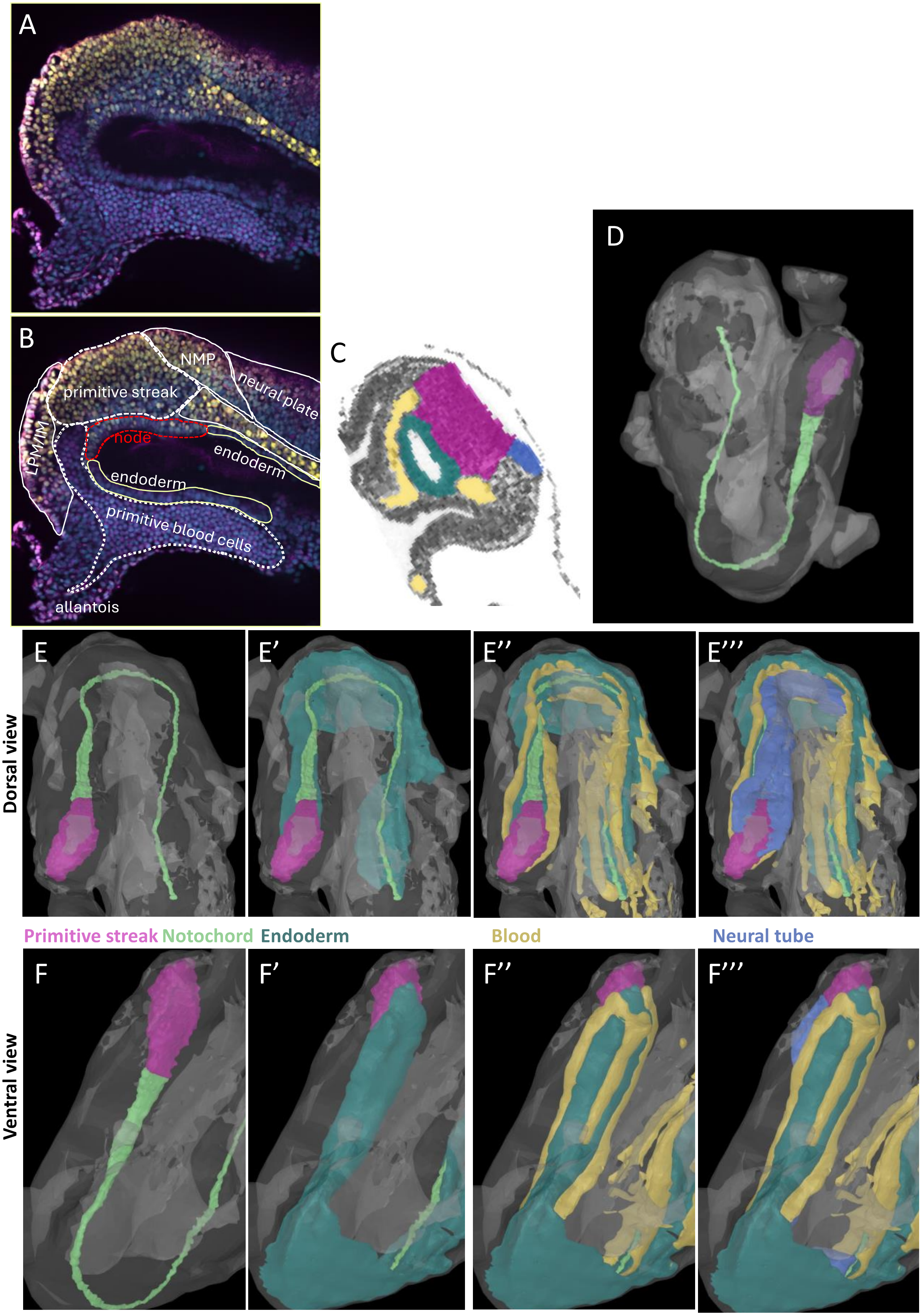
A
B
NMP
primitive streak
neural plate
LPM/IM
node
endoderm
endoderm
primitive blood cells
allantois
C
D
E
E'
E''
E'''
Dorsal view
Primitive streak
Notochord
Endoderm
Blood
Neural tube
F
F'
F''
F'''
Ventral view

**Supplementary Figure 4. *Wt1*- and *Bra*- expression domains in the mouse tail bud at E8.5.**
**A, B.** Images from Figure 3 (D, E) illustrating the different domains in optical transverse section of an E8.5 mouse embryo at primitive streak/node level.
**C.** Virtual section of an E8.5 (Theiler Stage 13) mouse embryo of the EMAP 3D Model (EMA:24), distance -104. Anatomy annotated via colour code: pink: primitive streak; dark green: endoderm; yellow: blood/blood vessels; purple: neural tube.
**D.** 3D Model embryo (EMA:24) showing notochord in light green and primitive streak in pink.
**E-E'''** and **F-F'''.** Dorsal view (E-E''') and ventral view (F-F''') of 3D Model embryo (EMA:24) showing primitive streak, notochord, endoderm, blood and neural tube (colour code as in C). Images C-F''' are screenshots obtained from the EMAP eMouse Atlas project website, eAtlasViewer (https://www.emouseatlas.org/eAtlasViewer_ema/application/ema/anatomy/EMA24.php). Data were accessed in July 2024.

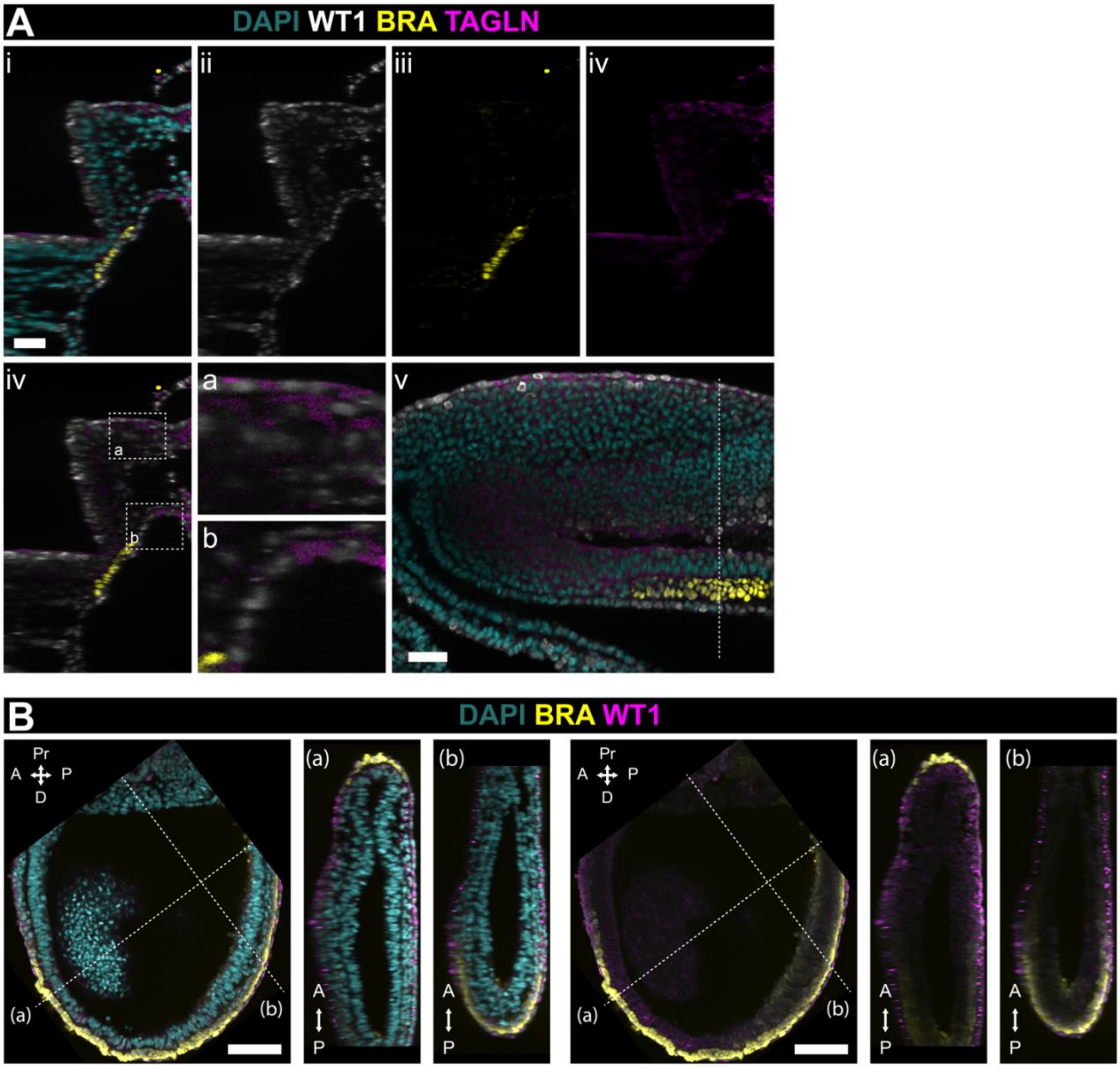


**Supplementary Figure 5. WT1 is co-expressed with TAGLN in the E8.5 mouse embryo, and expressed in the emerging mesoderm in the E7.5 mouse embryo.**

**A.** WT1 is co-expressed with BRA in the notochordal plate (i, ii, iii) and TAGLN in the mesoderm/LPM (iv, v). Oblique focal image in (vi) indicates focal plane shown in (i-v).

**B.** Whole mount staining of E7.5 embryo for BRA and WT1. Stipled lines indicate focal planes in (a) and (b).

Scale bars, 100 µm (B), 50 µm (A).

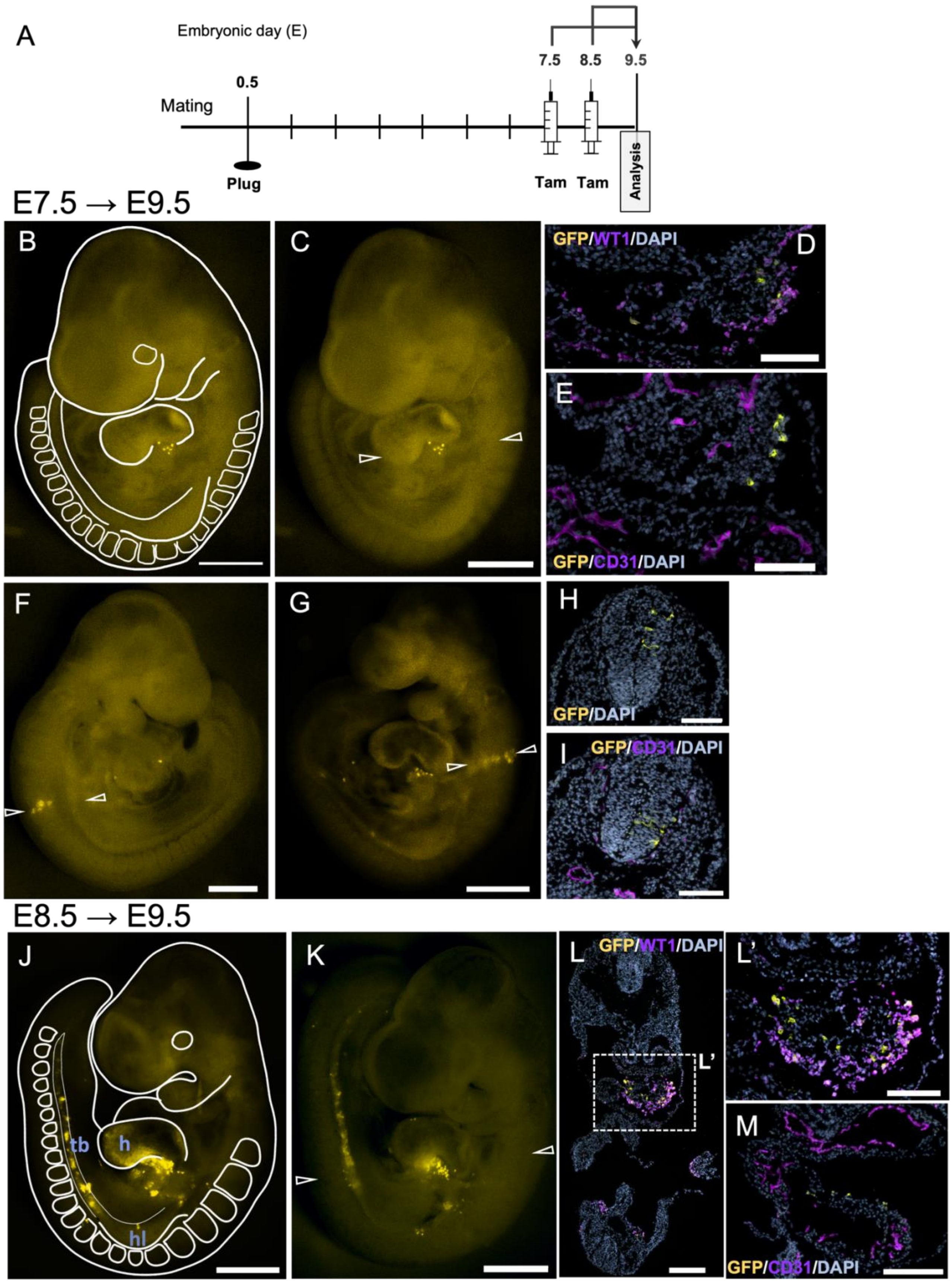


**Supplementary Figure 6.1.** ***Wt1*-lineage analysis at E9.5 after Tamoxifen dosing at E7.5 or E8.5.**

**A.** Schematic of the study design: CD1 dams were time mated with *Wt1*$^{CreERT2/+}$;*Rosa26*$^{mTmG/mTmG}$ males and received Tam at E7.5 or E8.5; embryos were dissected at E9.5.
**B, C.** Representative image of an E9.5 embryo after Tam dosing at E7.5 with schematic drawing overlay (**B**) highlighting the important developmental tissues. Very few GFP-positive cells were found within the pro-epicardium caudal to the heart tube. Arrowheads point to level of tissue sections for D, E.
**D, E.** Immunofluorescence staining of Wt1 and GFP (G) and CD31 and GFP (H) on sections showed a few GFP-positive cells within the epicardium and developing heart.
**F, G.** Representative images of E9.5 embryos after Tam dosing at E7.5 showing GFP-labelled cells within a distinct domain in the neural tube. Arrowheads point to level of tissue sections for H, I.
**H, I.** Immunofluorescence staining of GFP (H) and CD31 and GFP (I) on sections showed GFP-positive cells within the developing neuroepithelium.
**J.** Schematic drawing overlay over a representative E9.5 embryo after Tam dosing at E8.5.
**K.** Representative image of a second E9.5 embryo after Tam dosing at E8.5. GFP-positive cells were found within the pro-epicardium caudal to the heart tube, and in the urogenital ridge. Arrowheads point to level of tissue sections for L, L', M.
**L, L', M.** Immunofluorescence staining of Wt1 and GFP (D, D') and CD31 and GFP (E) on sections showed GFP-positive cells within the epicardium and developing heart and the urogenital ridge.
h, heart, hl, hindlimb, tb, tailbud, ur, urogenital ridge.
Scale bars, 500 µm (B, C, F, G, J, K), 200 µm (L'), 100 µm (D, E, H, I, L, M).

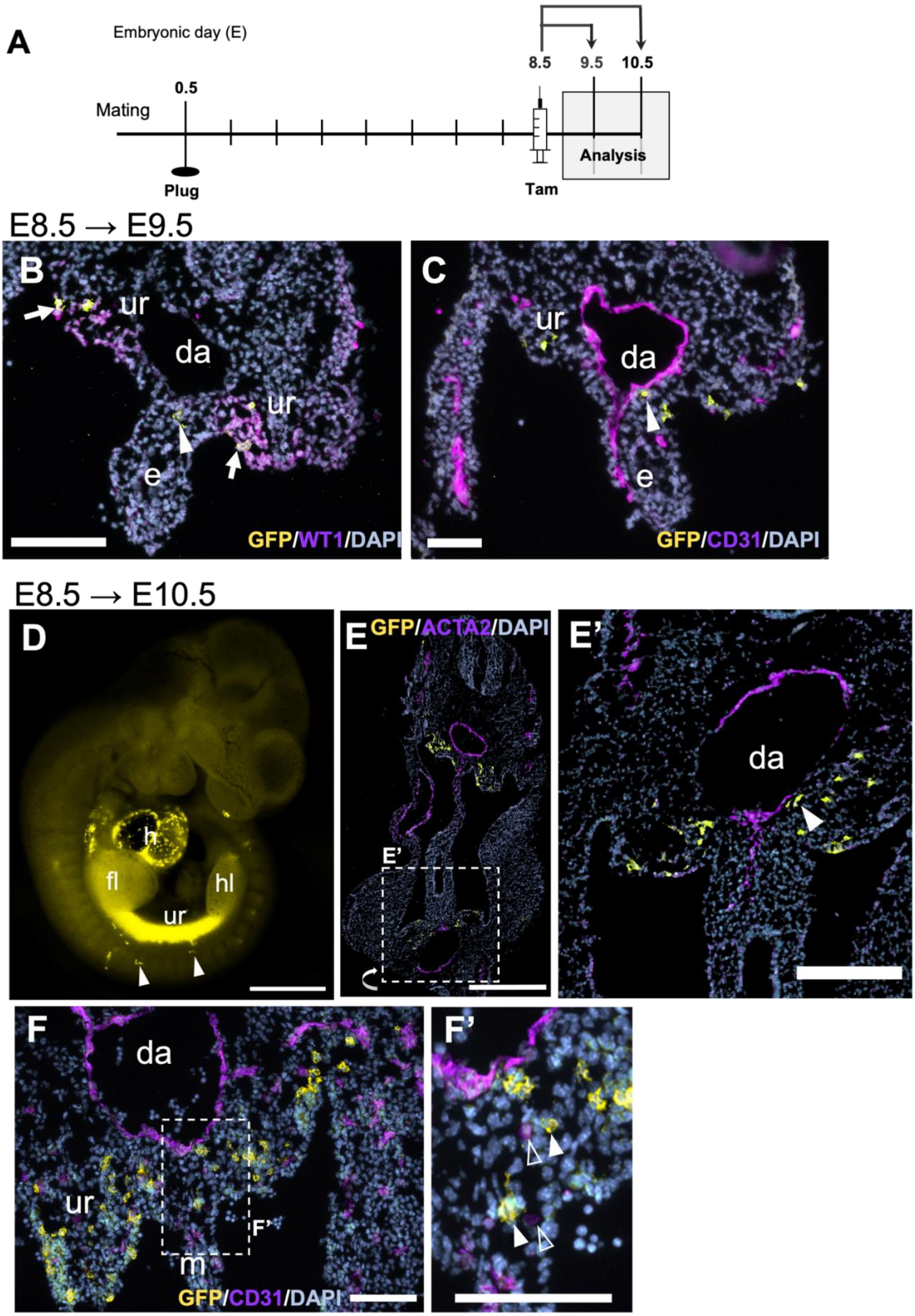


**Supplementary Figure 6.2. Localisation of *Wt1*-lineage cells in the urogenital ridge and close to the mesentery near endothelial cells.**

**A.** Schematic of the study design: CD1 dams were time mated with *Wt1*$^{CreERT2/+}$;*Rosa26*$^{mTmG/mTmG}$ males and received Tam at E8.5; embryos were dissected at E9.5 or E10.5.
**B, C.** Immunofluorescence staining for GFP and WT1 (**B**) and GFP and CD31 (**C**) on sections through E9.5 embryos after Tam dosing at E8.5. GFP-positive cells were detected in the region between dorsal aorta (da) and mesentery, besides presence in the urogenital ridge (ur).
**D.** Representative overview image of embryo at E10.5 after Tam dosing at E8.5. GFP-positive cells were most prominent in the heart (h), and urogenital ridge (ur). Arrowheads point to individual cells dorsally away from the main ur expression domain. The limb buds appeared brightly fluorescing, however this was based on the strongly fluorescing tissues (IM, heart) lying underneath.
**E, E'.** Immunofluorescence staining for GFP and ACTA2 on sections through the region just below the forelimb bud. GFP-positive cells were localized in the urogenital ridge and close to the dorsal aorta (da)(arrowheads).
**F, F'.** Immunofluorescence staining for GFP and CD31 on section through similar embryonic level as in E. GFP-positive cells were also found ventral to the dorsal aorta and dorsal to the mesentery, closely associated with cells of the endothelial plexus (filled arrowhead: GFP+, open arrowhead: CD31+).
da, dorsal aorta, e, endoderm, fl, forelimb bud, h, heart, hl, hindlimb bud, ur, urogenital ridge. Scale bars, 1000 μm (D), 500 μm (E), 200 μm (E'), 100 μm (B, C, F, F').

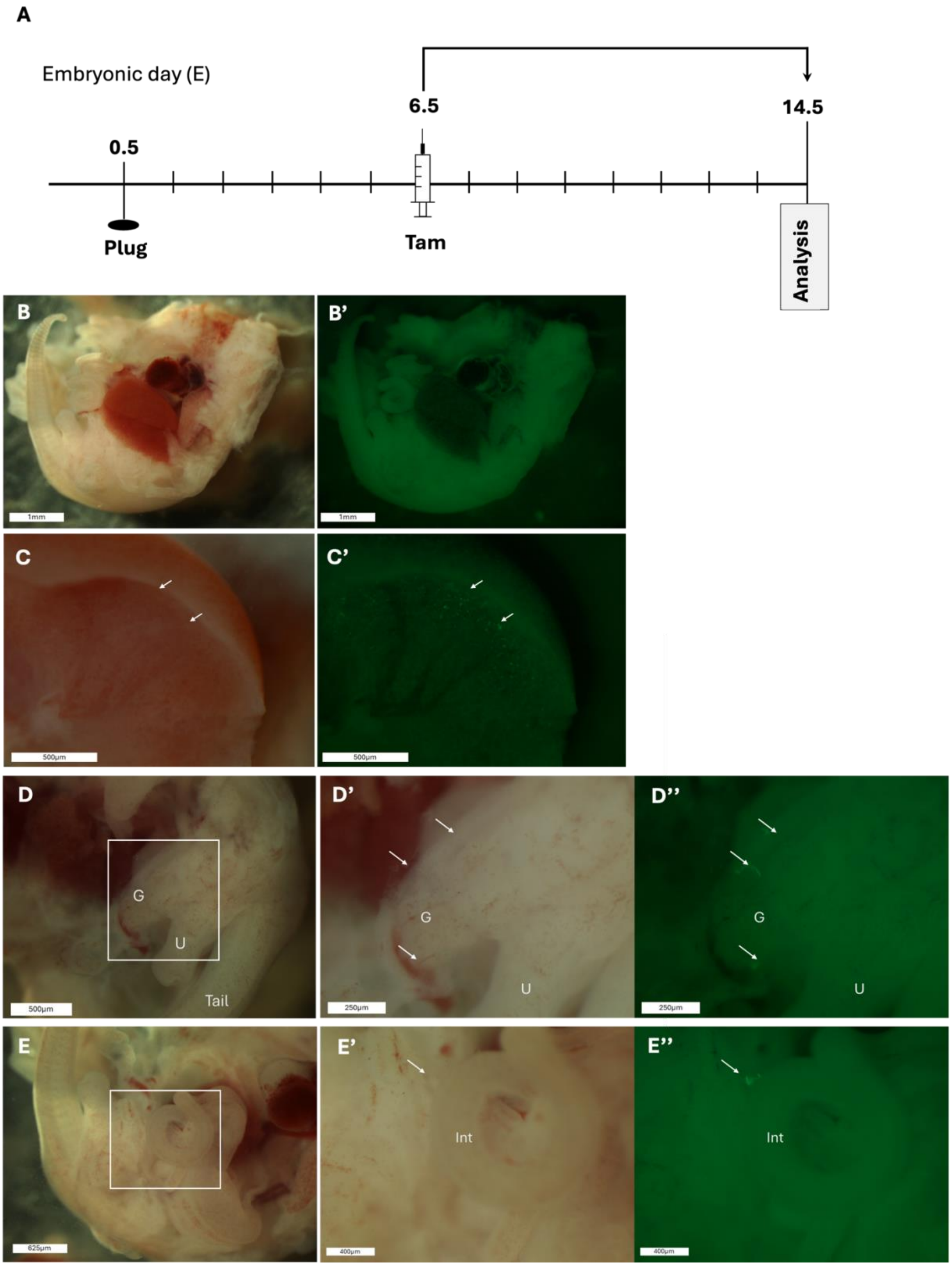


**Supplementary Figure 6.3. Lineage tracing of *Wt1* at E6.5 followed by end-point analysis at E14.5.**

**A.** Schematic of the study design: CD1 dams were time mated with $Wt1^{CreERT2/+}$;$Rosa26^{mTmG/mTmG}$ males and received Tam at E6.5; embryos were dissected at E14.5.

**B-B'.** Overview of embryo showing very few GFP+ cells.

**C-C'**. Inferior view of mesothelium lining the liver showing a few GFP+ cells (arrows).

**D-D”.** Caudal view showing the tail, urethra (U) and gonad (G) of the embryo with a few GFP+ cells on or near the intestine.
**E-E”.** Sagittal view focussed on the intestine (Int) of the embryo showing a GFP+ cell.

**Supplementary Table 2: Cell distribution in E9.5 *Wt1*$^{CreERT2/+}$;*Rosa26*$^{mTmG/+}$ embryos after Tam at E7.5.**

| Litter | Embryos | GFP-positive cells | | |
|---|---|---|---|---|
| | | Proepicardium | Neural tube | Trunk |
| 1 | 1 | 8 | 0 | 0 |
| | 2 | 6 | 0 | 4 |
| | 3 | 5 | 6 | 0 |
| | 4 | 6 | 0 | 0 |
| | 5 | 6 | 4 | 0 |
| 2 | 6 | 7 | 0 | 0 |
| | 7 | 10 | 3 | 5 |
| | 8 | 5 | 2 | 0 |
| | 9 | 8 | 4 | 3 |
| | 10 | 2 | 0 | 1 |
| 3 | 11 | 20 | 7 | 12 |
| | 12 | 12 | 0 | 7 |
| | 13 | 8 | 0 | 5 |
| | 14 | 6 | 0 | 11 |
| | 15 | 8 | 0 | 3 |

Analysis by counting cells in embryos using whole mount imaging (dissecting microscope with fluorescence capabilities).

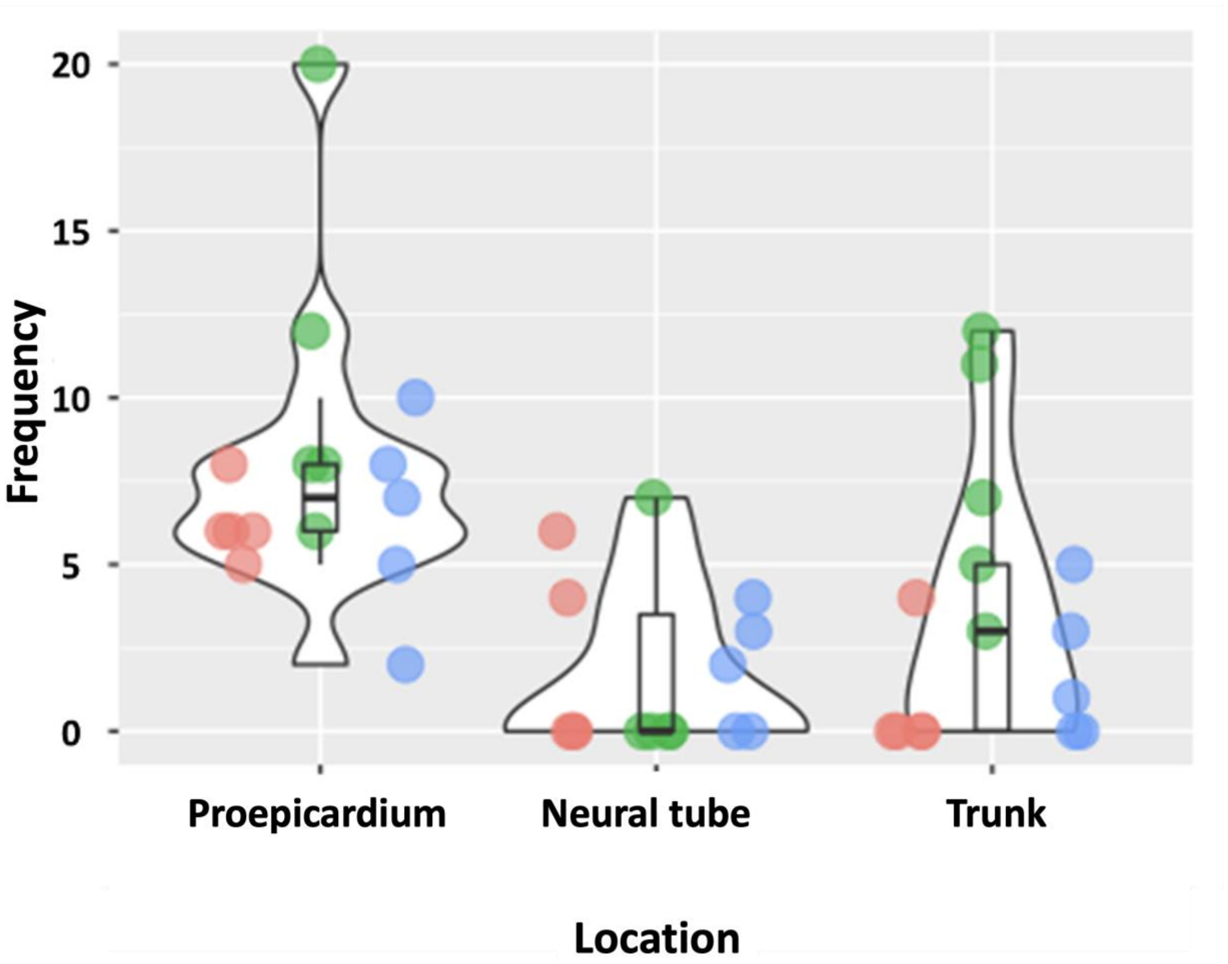


**Supplementary Figure 7. Violin plot displaying the distribution of GFP-positive cells found in three litters of E9.5 *Wt1*$^{CreERT2/+}$;*Rosa26*$^{mTmG/+}$ embryos after Tam at E7.5.** The GFP-positive cell population in the epicardium had a mean of 7 cells in the three litters. Numbers of GFP-positive cells in the neural tube had a mean of 4 cells in the three litters. Most of the embryos had 1 to 5 GFP-positive cells in the trunk area but others had up to 12 cells.

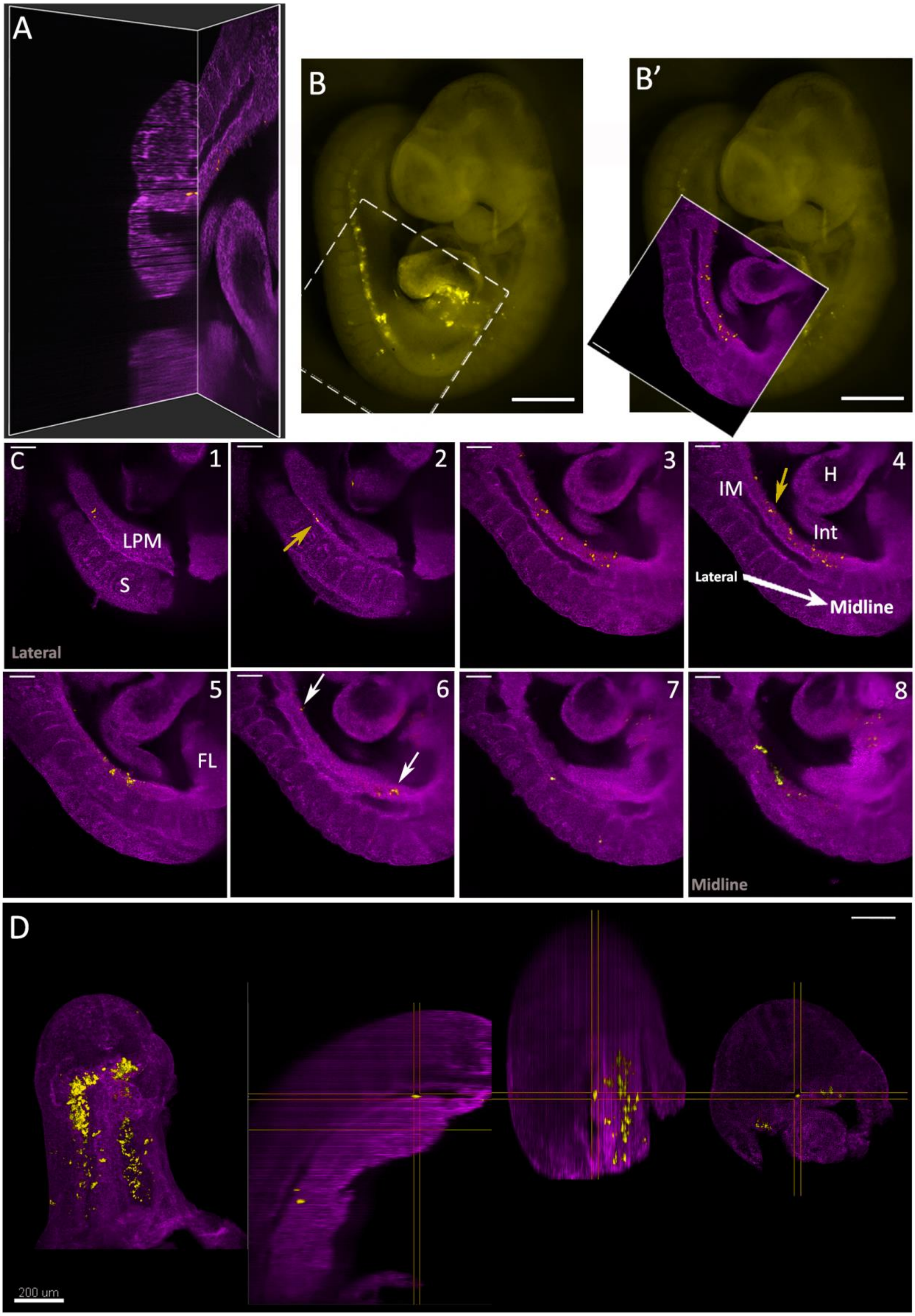


**Supplementary Figure 8: Light sheet microscopy analysis of Wt1 lineage at E9.5 after tamoxifen dosing at E8.5.**

**A.** GFP-positive cells (in yellow) were detected in the mesentery, located between the dorsal aorta and the intestine, using the IMARIS Ortho Slicer tool. The membrane-bound dTomato appears in purple to illustrate the body structure.

**B, B'.** Overview images combining dissecting microscopic and light sheet microscopic images to demonstrate the specific area of an embryo imaged in the light sheet microscope (LSM).

**C1-8.** Eleven selected sagittal slices from lateral side towards midline of the body from z-stacks after light sheet microscopy of the embryo shown in B. The images demonstrate patches of GFP-positive cells in the lateral plate and intermediate mesoderm (yellow arrows) as well as the tissue close to the dorsal aorta and mesentery (white arrows). LSM images were generated by the Slice View function in IMARIS software.

**D.** The posterior embryo region was imaged from different angles to visualise the GFP-positive cells located near the dorsal aorta wall. The size of the crosshair is 30 µm in all three axes X, Y and Z.

LPM, lateral plate mesoderm, IM, intermediate mesoderm, FL, forelimb, Int, intestine, S, somites. Scale bars, 500 µm (B, B', for DM), 200 µm (A, C1-8, D for LSM).

**Supplementary Movie:**
https://drive.google.com/file/d/1PlzNmE2lt2-IJXVNeGhpNe_Prs5pgCTQ/view?usp=share_link

The 3D animation was generated from z-stacks obtained with light sheet microscopy of an E9.5 *Wt1*$^{CreERT2/+}$*;Rosa26*$^{mTmG/+}$ embryo after Tam at E8.5. The membrane-bound dTomato appears in red (561 nm) to illustrate the body structure and the GFP-positive cells appear in green.

GFP-positive cells were labelled using the Imaris software according to their location:
in the mesentery – white spots;
in the proepicardium and within the trunk area – blue spots;
in the neural tube – yellow spots;
in the lateral plate mesoderm – green spots.

**Supplementary Table 3: The number of GFP-positive cells counted in light-sheet microscopy images of *Wt1$^{CreERT2/+}$;Rosa26$^{mTmG/+}$* embryos at E9.5 after Tam administration at E8.5.**

| Embryos analysed | GFP-positive cells in the trunk | GFP-positive cells in the epicardium | GFP-positive cells in the NT | Total number of GFP-positive cells |
|---|---|---|---|---|
| 1 | 43 | Not included (*) | 3 | 46 |
| 2 | 28 | Not included (*) | 2 | 30 |
| 3 | 37 | 7 | 2 | 46 |

The number of positive GFP-positive cells in the trunk were between 28 and 43 cells. Only one sample contained the heart area and showed about seven cells in the epicardium. The GFP-expression in the neural tube was restricted to only two to three cells in the analysed embryos.
*(*) The embryos were only imaged in the area of the trunk and the heart region had not been included.*
